# Revealing function-altering MECP2 mutations in individuals with autism spectrum disorder using yeast and *Drosophila*

**DOI:** 10.1101/2024.08.13.607763

**Authors:** Eric Chen, Jessica Schmitt, Graeme McIntosh, Barry P. Young, Tianshun Lian, Jie Liu, Kexin K. Chen, J. Beatrice Liston, Lily MacDonald, Bill Wang, Sonia Medina Giro, Benjamin Boehme, Mriga Das, Seevasant Indran, Jesse T. Chao, Sanja Rogic, Paul Pavlidis, Douglas W. Allan, Christopher J.R. Loewen

## Abstract

Pathogenic variants in MECP2 commonly lead to Rett syndrome, where MECP2’s function as a DNA cytosine methylation reader is believed critical. MECP2 variants are also catalogued in individuals with autism spectrum disorder (ASD), including nine missense variants which had no known clinical significance at the start of this study. To assess these nine variants as risk alleles for ASD, we developed MECP2 variant functional assays using budding yeast and *Drosophila*. We calibrated these assays with known pathogenic and benign variants. Our data predict that four ASD variants are loss of function and five are functional. Protein destabilization offers insight into the altered function of some of these variants. Notably, yeast and *Drosophila* lack DNA methylation, yet all Rett pathogenic and ASD variants located in the methyl DNA binding domain that we analyzed proved to be loss of function, suggesting a clinically-relevant role for non-methyl DNA-binding by MECP2.

## INTRODUCTION

The X-linked protein MECP2 was first identified in screens for proteins recruited to methylated DNA (Lewis et al. 1992), and loss of function (LoF) alleles of MECP2 were subsequently found as causative in the neurodevelopmental disorder, Rett syndrome (Amir et al. 1999). Since then variants in MECP2 have been found in most cases of Rett, but also in other neurodevelopmental disorders, including autism spectrum disorder (ASD) (Wan et al. 1999; Amir et al. 2000; Nomura 2005; Chahrour and Zoghbi 2007). The origin and phenotypic consequence of MECP2-damaging variants differ notably between sexes. In females, the vast majority of these variants are *de novo*, while in males, they can also be inherited from unaffected mothers (Ananth et al. 2024). Variants that typically result in Rett syndrome in females tend in males to cause severe neonatal encephalopathy, often leading to death in the first year of life. However, in males with two populations of X chromosomes, such as those with Klinefelter syndrome (47XXY) or somatic mosaicism, these variants result in a milder Rett-like condition known as male RTT encephalopathy. Males can also be affected by variants that are seemingly benign in females, which can lead to moderate to severe intellectual disability (Pascual-Alonso et al. 2021).

MECP2 has largely been examined as a DNA cytosine methylation reader via its methyl cytosine binding domain (MBD) that recruits co-repressor complexes to DNA to repress transcription via its transcriptional repressor domain (TRD) (Jones et al. 1998; Ng and Bird 1999; Lunyak et al. 2002; Harikrishnan et al. 2005). The MBD, and to a lesser extent the TRD, harbors most Rett LoF variants (Ehrhart et al. 2021), resulting in variants which are believed to reduce binding to methylated DNA or reduce recruitment of co-regulator complexes, respectively (Ballestar et al. 2000; Yusufzai and Wolffe 2000). While numerous MECP2 variants have been found in patients that exhibit Rett with ASD, recent reports have annotated five MECP2 missense variants in ASD without an associated Rett diagnosis (R91W, G195S, A202G, E282G, S411R) (Schaaf et al. 2011; Wang et al. 2016; Wen et al. 2017; Guo et al. 2018; Leblond et al. 2021). However, it is unclear whether any of these variants are function altering and clinically significant. Two of them, R91W and A202G, are classified by ClinVar (Landrum et al. 2018) as variants of uncertain significance (VUS) in cases of severe neonatal onset encephalopathy with microcephaly. E282G is classified by ClinVar as likely pathogenic. The remaining two variants (G195S and S411R) have no current ClinVar classification.

While MECP2 is understood to primarily be a methyl-cytosine DNA (DNAm) binding protein, it has become clear it also binds unmethylated DNA with a preference for GC-rich sequences (Georgel et al. 2003; Hansen et al. 2010; Skene et al. 2010; Cheng et al. 2014). This has been proposed as an alternate hypothesis for why MECP2 appears to preferentially bind methylated GC-rich regions of the genome, as opposed to binding DNAm methylation *per se*. It can be challenging to segregate DNAm-dependent from DNAm-independent functions of MeCP2 *in vivo*, as DNAm is both abundant and essential in mammals. Consequently, it is still unclear what fraction of MeCP2’s functions are dependent on its interaction with methylated and/or unmethylated DNA. Such ambiguity leaves unresolved the question of whether disease relevant variants in MECP2 selectively disrupt binding to methylated versus unmethylated DNA.

There are extremely low levels of methylated cytosine in the *Drosophila* genome, and none in *Saccharomyces cerevisiae* (Lyko et al. 2000; Buitrago et al. 2021). Thus, fly and yeast offer avenues to explore the interactions of MECP2 with unmethylated DNA, and to test if Rett and ASD variants alter MECP2 binding to unmethylated DNA. Moreover, while neither organism has an ortholog of *MECP2*, both organisms are robust genetic models that have proven well suited to comparing the function of large numbers of gene variants (Post et al. 2020; Young et al. 2020; Ganguly et al. 2021; Her et al. 2024). MECP2 has been expressed in *Drosophila*, where it associates with chromatin and mediates transcriptional repression via interactions with orthologs of known MECP2 physical interactors in humans, including Sin3A, N-CoR, REST as well as other chromatin modifiers (Cukier et al. 2008). In fly neurons, MECP2 impacts dendritic branching and behavioural output (Vonhoff et al. 2012; Williams, White, et al. 2016) and C-terminal truncation mutants (R294X specifically) promote neuronal apoptosis (Williams, Mehler, et al. 2016). In yeast, MECP2 has recently been shown to bind chromatin and mediate widespread transcriptional repression (Brown et al. 2025 Mar 18).

Our goal in this study was to use the *Drosophila* and budding yeast models to test the functional impact of these ASD MECP2 variants to determine whether they may confer autism risk, by virtue of their altered function. We calibrated our assays with MECP2 variants classified by ClinVar as pathogenic or benign, in accordance with American College of Medical Genetics and Genomics (ACMG) guidelines for experimental assays (Brnich et al. 2019); first to validate our assays for variant assessment, and second to provide a rubric for determining if test ASD variants are likely pathogenic or likely benign. All calibration variants functioned as expected in the *Drosophila* assay, indicating that we could functionally assess any test variants. The yeast assay could be calibrated appropriately for the MBD domain, but the MBD proved to be the only domain showing the expected separation between benign and pathogenic calibration variants.

Our analyses indicate four of the nine test ASD variants were LoF, making these variants candidate risk alleles for ASD. Interestingly, these variants mostly proved to be less severe LoF than Rett-associated pathogenics, providing a possible explanation for how MECP2 mutations can lead to ASD, but not Rett.

## METHODS

### MECP2 isoform selection

*MECP2* has two major isoforms, E1 and E2. We selected to work with *MECP2* isoform E2 (ENST00000303391, NM_004992) based on the recommendation by the ClinGen Rett/Angelman-like Expert Panel (McKnight et al. 2022). This transcript is annotated as the APPRIS (Annotation of Principal and Alternative Splice Isoforms (Rodriguez et al. 2013)) principal isoform in Ensembl genome browser, based on the annotations for protein structure, function and cross-species conservation, and is the highest expressed transcript in the brain (GTEx portal; gtexportal.org).

### Variant aggregation, annotation and prioritization

We utilized our in-house computational pipeline to aggregate *MECP2-E2* missense variants from ClinVar and VariCarta and perform comprehensive annotation. We collected ClinVar variants (downloaded in January 2022) that have at least one gold review star, i.e., the review status (CLNREVSTAT) has to have one of the following designations: “criteria provided, single submitter”, “criteria provided, conflicting interpretations”, “criteria provided, multiple submitters, no conflicts” or “reviewed by expert panel”. For each of these variants, we also obtain its interpretation (clinical significance; CLNSIG) and reported condition (disease name; CLNDN). The ASD missense variants were obtained from VariCarta, a database of variants found in individuals diagnosed with ASD and reported in the scientific literature, that we created (Belmadani et al. 2019). We annotated the aggregated variants using the Ensembl VEP tool (McLaren et al. 2016). We also added protein domain annotation relevant in the context of Rett Syndrome (Good et al. 2021).

### MECP2-E2 cloning

A *MECP2-E2* cDNA clone was obtained (Horizon, Clone Id: 3956518, # MHS6278-202830559), and *MECP2* coding sequences were amplified with iProof DNA polymerase (Bio-Rad, California USA, #1725301) using the following primers: GA-attL1-MECP2-522-F (tgtacaaaaaagcaggctccggcaccatggtagctgggatgttag) GA-attL1-MECP2-522-R (ttgtacaagaaagctgggtctatcagctaactctctcggtc) Amplicons were cloned in PCR8-GW-TOPO (Invitrogen, ThermoFisher Scientific, Waltham, USA, catalog #K250020) using NEBuilder HiFi DNA Assembly Master Mix (New England Biolabs, USA, # M5520). The resulting construct was named MECP2-pENTRY. A sequence of CCGGCACC was introduced in primer GA-attL1-MECP2-522-F (underlined) preceding the start codon for *MECP2-E2*. After LR cloning of MECP2-pEntry into pDEST vectors, this CCGGCACC sequence can function as a Kozak sequence to initiate translation, or as an in-frame linker for N-terminal peptide sequences. Missense variants were generated in MECP2-pENTRY either using the Q5 mutagenesis kit (New England Biolabs, USA, # E0554) in house, or were made by GenScript (New Jersey, USA). For MECP2 expression clone construction, MECP2-pENTRY plasmids were subcloned in pAG426GAL-ccdB (a kind gift from a gift from Susan Lindquist; Addgene plasmid # 14155) or pGW-HA.attB (Bischof et al. 2007) using the Gateway LR Clonase reaction (Thermo Fisher Scientific, Waltham, USA, catalog # 11791020). All transgenic *UAS-MECP2* variants (on pGW-HA.attB) *Drosophila* strains used were verified by Sanger sequencing (UBC NAPS, Vancouver, Canada) or whole amplicon sequencing (Plasmidsaurus, CA, USA) (Supplemental Data 1).

### Sentinel interaction mapping (SIM) in yeast

SIM screening for *MECP2* was performed as described previously (Young et al. 2020). A singer RoToR HDA colony arraying robot was used to copy arrays of yeast cells. All steps were performed at a density of 1,536 colonies per plate and plates were incubated for 24 hours at 30°C prior to the next pinning step, except where indicated otherwise. The SGA query yeast strain Y7092 (Tong et al. 2001) was transformed with plasmid pSI7338-MECP2 and arrayed on synthetic defined medium lacking uracil. The haploid yeast deletion mutant array (DMA) was arrayed across a total of four plates on YPD medium containing 200 mg/L G418 sulfate (YPD+G418). The query array was mated to the DMA by subsequently pinning both arrays on to fresh YPD plates. Diploids were selected by pinning colonies to synthetic defined medium lacking uracil containing 200 mg/L of G418 sulfate and subsequently pinned to YPD+G418 plates. Each plate was then pinned in triplicate to enriched sporulation media (Tong et al. 2001) and incubated at 22 °C for seven days. Following this, *MAT*a haploid cells were germinated by pinning to synthetic complete medium lacking histidine, arginine and lysine with the addition of 50 mg/L canavanine sulfate, 50 mg/L thialysine and incubating for two days. To select for deletion strains carrying the MECP2-Ref plasmid, colonies were pinned to two pairs of plates (“control” and “experimental”) containing synthetic complete medium lacking histidine, arginine, lysine and uracil, containing 50 mg/L canavnine sulfate, 50 mg/L thialysine, 200 mg/L G418 sulfate (“HURK+G418” medium). The experimental set was supplemented with 10 nM - estradiol to induce expression of *MECP2*; this was omitted from the control set. The final plates used for imaging were generated by copying each of these plates once more to the same medium. Plates were scanned using a CanoScan 8600F flatbed scanner and subsequently analyzed using *Balony* software (Young and Loewen 2013) to compare the normalized colony size for each strain in the presence or absence of estradiol. Genetic interactions were defined as those strains where the ratio of experimental:control colony size was below a threshold determined by *Balony* in all three replicates and with a p-value < 0.05. The strains *pho85Δ, rad50Δ, slx5Δ, swi4Δ, swi6Δ,* and *xrs2Δ* were confirmed to be correct by PCR amplification of down-tag barcodes with the standard primers B-U1 and B-D1 and sequencing.

### Yeast quantitative liquid growth assays

Yeast strain JHY716 was transformed with various plasmids containing *MECP2* variants placed downstream of the *GAL1* promoter. Four independent clones were picked from transformation plates and transferred to SD-URA media containing 2% raffinose, 2% galactose to induce expression of *MECP2*. For each clone, a 2 mL culture in SD-URA with 2% raffinose, 2% galactose was grown overnight at 30 °C in a shaking incubator. The following morning, cultures were diluted to an OD600 of ∼0.25 in the same media. After 6 hours of incubation at 30 °C, log phase cultures were diluted to an OD600 of 0.125. Three aliquots of 200 µl were taken from each culture and transferred to a well of a flat-bottomed 96-well dish. Growth of cultures was measured overnight at 30 °C with shaking at 800 rpm in an Agilent Logphase 600 plate reader.

All plates contained strains expressing wild type *MECP2* and the corresponding empty vector as controls. The exponential growth constant k was determined for each well by fitting to an exponential growth equation during the exponential phase (typically between 5 and 10 hours post dilution).

### Protein abundance assays

To determine the abundance of MECP2 variants in yeast, strains were grown in SD-URA media containing 2% raffinose to early log phase. MECP2 expression was induced by the addition of 2% galactose to the growth medium. Cells were harvested three hours later by centrifugation and stored at -80 °C for subsequent processing. Cell pellets were resuspended in Laemmli sample buffer, and heated to 95 C for 5 minutes. Cell lysis was achieved by the addition of 50 µl acid-washed glass beads followed by two rounds of disruption in a MP Biomedical FastPrep-24 homogenizer (maximum speed for 60 s with 5 minutes cooling between cycles). Samples (0.2 OD600 equivalents) were separated by SDS-PAGE on Bio-Rad 4-15% pre-cast gels and transferred to nitrocellulose membranes. MECP2 was immunoblotted using Sigma antibody (M9317), with PGK1 (Abcam AB113687) used as a loading control. For visualization, secondary antibodies conjugated to Cy3 (MECP2) and AlexaFluor-488 (PGK1) were used and imaged using a Bio-Rad ChemiDoc MP Imaging System. MECP2 band volume was normalized to PGK1 band volume for each variant.

### Drosophila strains

All flies were maintained on standard cornmeal food at 25 °C and 70% humidity unless otherwise specified. *P{GAL4}A9* (BL8761) (referred to as *A9-GAL4*) was obtained from Bloomington *Drosophila* Stock Centre (Indiana, USA). *UAS-MECP2* reference (*UAS-MECP2-ref*) and variant sequences (collectively termed *UAS-MECP2.#*) were integrated into embryos of genotype *y[1] w[] P{y[+7.7]=nos-phiC31\int.NLS}X; P{y[+t7.7]=CaryP}attP40* (referred to as attP40) by Genome ProLab (Quebec, Canada), and maintained as *w[1118]; P{y[+t7.7] w[+mC]=UAS-MECP2.#.GW.\**}*attP40*, where # represents the variant and * represents integrant line number (Supplemental Data 1).

### Crossing scheme

Experimenters were double blinded to genotype throughout. Homozygous *UAS-MECP2* males and a control *attP40* line were crossed to *A9-GAL4* virgins (ratio 4 males : 12 females), and maintained at room temperature (∼21 °C) for 72 hours. Crosses were then transferred into fresh ‘assay’ vials, and transferred each 24 hrs each for three days, generating three ‘assay’ vials per cross. After removal of adults from each vial, vials were placed into a 29 °C incubator (70% humidity). Three days after eclosion, adult flies were sorted by sex and placed in pure isopropanol at -20 °C. All progeny flies for comparison were tested in a single batch.

### Wing imaging

Experimenters were double blinded to genotype throughout. Flies were removed from isopropanol, dried on paper towels, and 10-25 left wings were removed per genotype per assay vial (one wing per fly). Wings were mounted on microscope slides in Canadian Balsam Fir (1.01691.0025, CAS-No: 8007-47-4, Sigma Aldrich, MA, USA), coverslipped and sealed with nail polish. Females have larger wings and the X-linked *A9-GAL4* driver is expressed at higher levels in males, due to X-chromosome male-hyperactivation. Thus, it is important to analyze males and females separately. While the impact of MECP2 variants in the wing was similar between males and females, the phenotype was substantially stronger in males due to the higher GAL4 levels; hence, higher MECP2 levels. To simplify analysis, we therefore only fully analyzed male wings. Slides were compressed while drying for 24 hrs. Mounted wings were scanned using a Panoramic Midi II digital slide scanner (3DHISTECH, Budapest, Hungary).

Scanned fly wing images were cropped at 15x zoom using CaseViewer (3DHISTECH, Budapest,Hungary). Images were imported into Imaris v10.0 software (Bitplane, MA, USA) for measurement of wing vein area per wing area, using Labkit machine learning and Fiji software (University of Wisconsin-Madison, USA). In Labkit, we developed “BIGSMOOTHCLASSYBOI”, a machine learning algorithm to automate measurement of wing vein area for calculation of wing vein area/wing area (details provided in Supplemental Methods 1).

### Statistical analysis of variant effects

Yeast growth constants (k) and fly relative wing vein area data were analyzed using methods similar to Post et al. (Post et al. 2020). Briefly, linear mixed-effects models were fit to each data set using the *lmerTest* R package (Kuznetsova et al. 2017) to estimate the phenotypic effects of each variant compared to the *MECP2-ref* constructs, treating experimental batches as random effects (for R scripts see Supplemental Methods 2 and 3). For visualization, values were standardized to the range [0-1] where 0 represents the mean phenotype measured for an empty vector, and 1 represents the mean phenotype with the *MECP2-ref* construct. Four biological replicates, each with three technical replicates were performed for yeast variant growth assays, while for *Drosophila* wing assays 10-25 wings, each from a different animal were imaged per genotype.

## RESULTS

### Selection of variants

We selected benign and pathogenic variants with clear clinical classifications to calibrate our functional assays, and nine test (ASD) variants (Figure 1, Supplemental Data 1). We focused on ClinVar variants with “pathogenic”, “likely_pathogenic”, “benign” and “likely_benign” clinical interpretation as calibration variants. Priority was given to those with either “reviewed by expert panel” or “criteria provided, multiple submitters” review status. A few exceptions were made.

**Figure 1:**
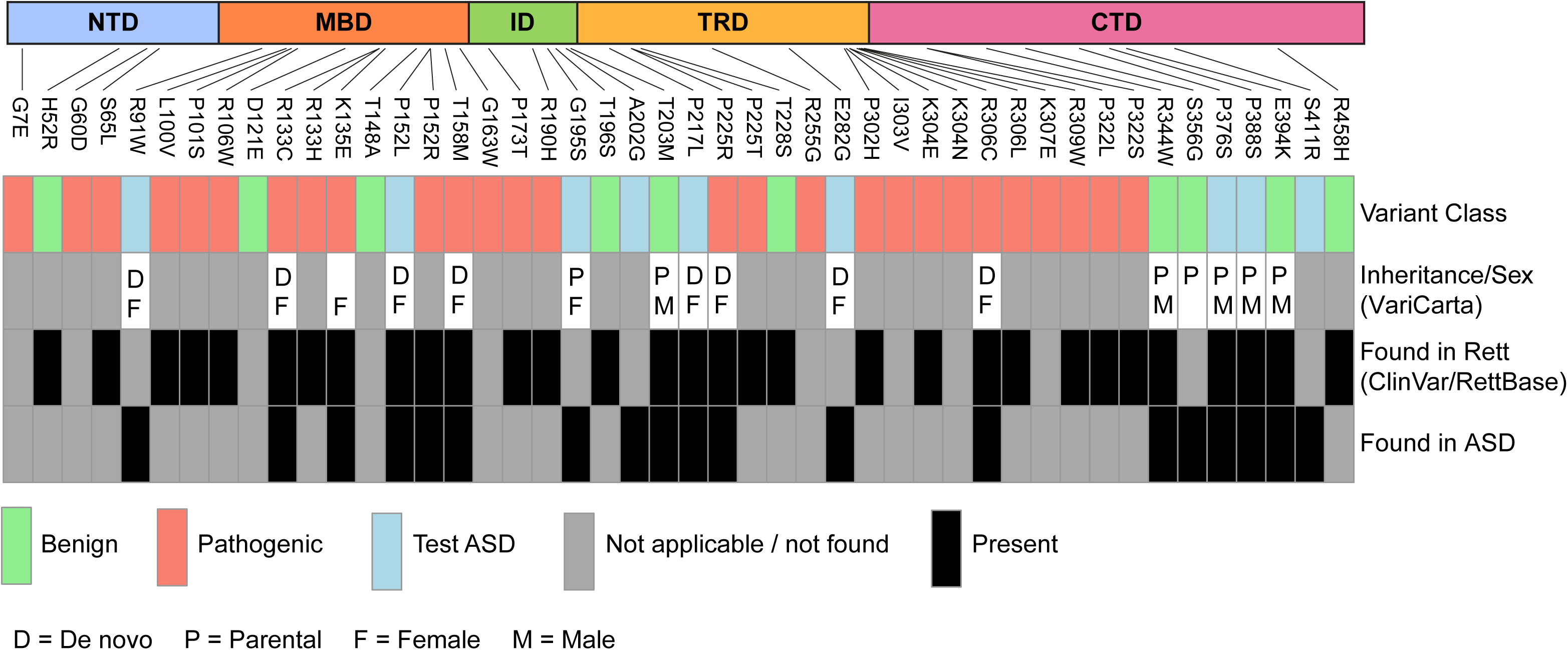
The forty-six MECP2 variants examined in this study. Domain locations of variants as well as variant class, variant inheritance, and whether they are found in Rett Syndrome and/or ASD. NTD – N-terminal domain, MBD – Methyl-CpG-binding domain, ID – Interdomain, TRD – Transcription repressor domain, CTD – C-terminal domain.

D121E, a single-submitter benign variant, and T148A, originally classified as benign, but now listed as conflicting were selected because there were no other benign variants in the MBD that matched the aforementioned criteria. When selecting pathogenic variants, in addition to using review status as a criterion, we also prioritized the ones with “Rett syndrome” condition annotation as well as the ones that have been frequently observed in Rett patients (Good et al. 2021). In total, we selected 37 calibration variants, 10 benign and 27 pathogenic. We selected ASD MECP2 variants from VariCarta as test variants as long as they did not have a clear clinical interpretation in ClinVar, for the total of 9 test (ASD) variants. Since the time of our initial selection, six test (ASD) variants have been given updated classifications in ClinVar: P152L, P217L and E282G becoming pathogenic/ likely pathogenic, and G195S, P376S and P388S becoming benign (Figure 1 displays the initial classifications).

### MECP2 produces a slow growth phenotype in yeast

*MECP2-ref* was cloned into a high-copy 2-micron plasmid designed to give controlled expression in response to addition of estradiol. Using an anti-MECP2 antibody we monitored MECP2 protein levels by Western blot in response to increasing estradiol (Figure 2A). In the no-estradiol condition we detected low levels of MECP2 protein, which increased with increasing estradiol and then saturated at ∼10 nM. Next, we measured the growth rate of WT yeast containing either the *MECP2-ref* plasmid or an empty vector control in response to estradiol addition (Figure 2B). Compared to empty vector which had little effect in response to increasing estradiol, yeast containing MECP2 showed a dose-dependent decrease in exponential growth rate. We compared this to another vector designed for strong overexpression utilizing the *Gal1/10* promoter in a 2-micron vector and observed an even greater decrease in growth rate in the presence of MECP2 (Figure 2C). Thus, expression of MECP2 reduced the fitness of WT yeast in a dose-dependent manner.

**Figure 2:**
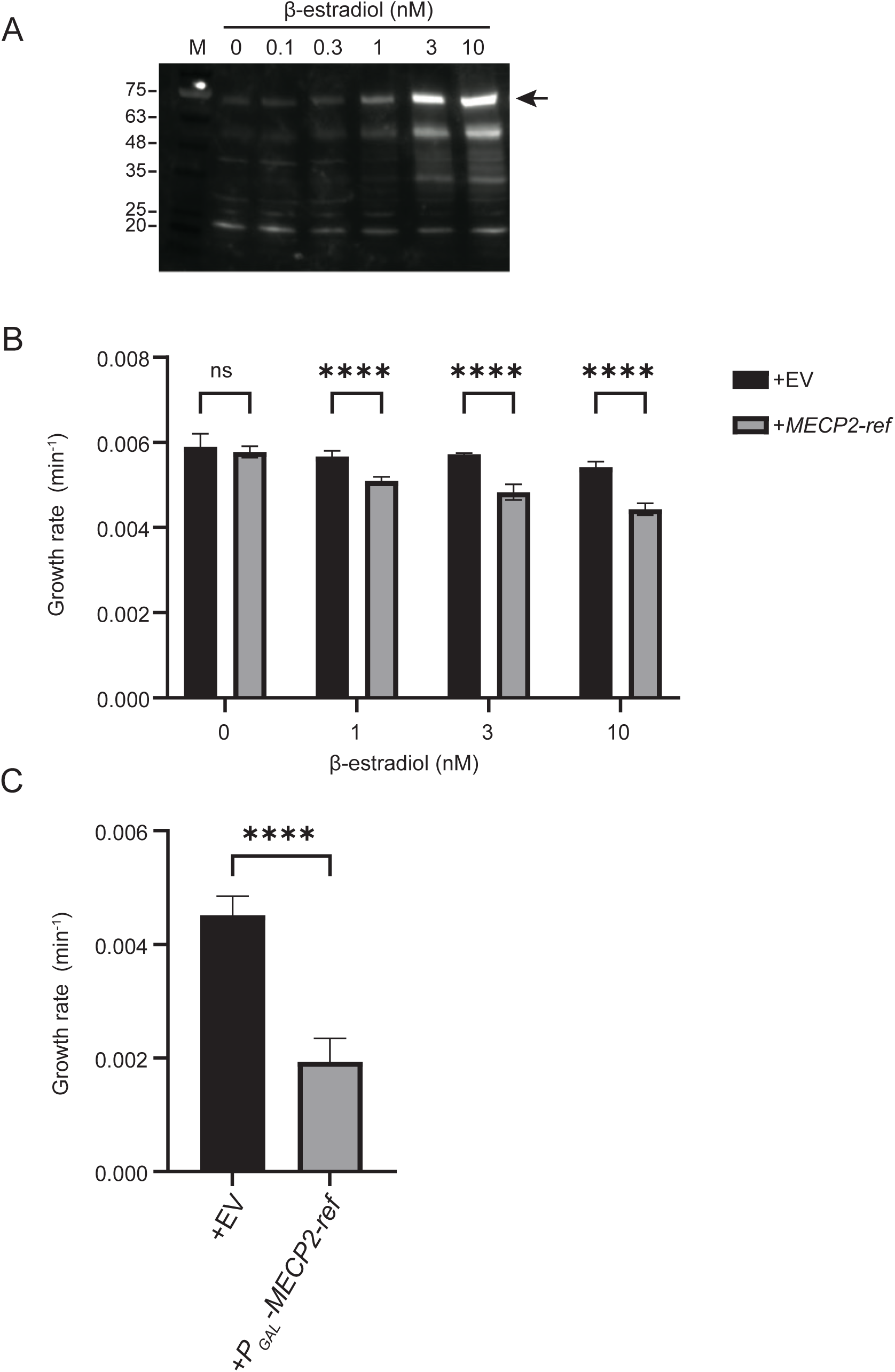
MECP2 expression and effect on growth in yeast. (A) Untagged MECP2-ref expressed from a 2-micron plasmid utilizing the estradiol-responsive promoter at various estradiol concentrations in WT yeast. Equivalent amounts of yeast were loaded into each lane and western blotted with anti-MECP2 antibody. M – molecular weight markers, arrowhead – full length MECP2. (B) Effect of MECP2-ref expression on growth of WT yeast from a plasmid utilizing the estradiol-responsive promoter at various estradiol concentrations. +EV – empty vector in wild type yeast, +MECP2-ref – MECP2-ref plasmid in wild type yeast. (C) Similar to B except using a 2-micron plasmid containing a galactose-responsive promoter and grown in galactose. **** – indicates p value < 0.0001. ns – not significant.

### MECP2 expression affects DNA-related processes in yeast

To understand how MECP2 might be functioning in yeast and if this is related to the normal function of MECP2 we employed a genetic interaction mapping approach we previously developed to study the function of human genes in yeast, called Sentinel Interaction Mapping (SIM) (Young et al. 2020). For SIM we express the human gene of interest in a collection of nearly 4,800 yeast deletion mutants and systematically identify genetic interactions, which indicate functional associations between the human gene and yeast cellular pathways. Hence, these genetic interactions will inform us on the function of the human gene in yeast and if this is relevant to its function in humans. We performed a SIM screen with *MECP2-ref* in which MECP2, under control of the estradiol responsive promoter in the 2-micron vector, was expressed in ∼ 4,800 different yeast deletion mutant strains in high-density arrays (see Methods for details). Colony sizes were scored using Balony software and genetic interactions were identified by comparing control arrays containing no estradiol (uninduced) to experimental arrays containing 10 nM estradiol (induced). We chose 10 nM estradiol because MECP2 protein expression saturated at this concentration and the effect on growth was not too severe, which would otherwise hinder identification of genetic interactions.

The SIM screen identified 123 aggravating genetic interactions in 3/3 biological replicates (Supplemental Data 2). We examined functional enrichment for these interactions using Gene Ontology (GO) analysis (Figure 3A; also see Supplemental Data 3 for gene lists). This revealed a network of interconnected functions related to DNA and chromosome stability, with the most significant enrichment being for the terms “Cellular response to stress” (30 genes) and “DNA damage response” (24 genes). Functions related to DNA binding (27 genes), cell cycle (21 genes), Golgi (8 genes) and nitrogen compound utilization (17 genes) were also enriched for. The large enrichment for DNA-related functions was consistent with MECP2 being a DNA binding protein with roles in transcriptional repression and chromatin organization. We focused on a subset of interactions from GO terms most relevant to MECP2 function. Shown in Figure 3B is the effect on growth of the individual mutants from the screen upon MECP2 induction for the GO terms “Chromatin organization”, “DNA binding” and “DNA damage response”. Because these are aggravating interactions, MECP2 induction (10 nM estradiol) resulted in decreased growth rate of the mutants relative to no induction (0 nM estradiol). Growth was normalized to account for the effect of MECP2 induction on WT yeast growth.

**Figure 3:**
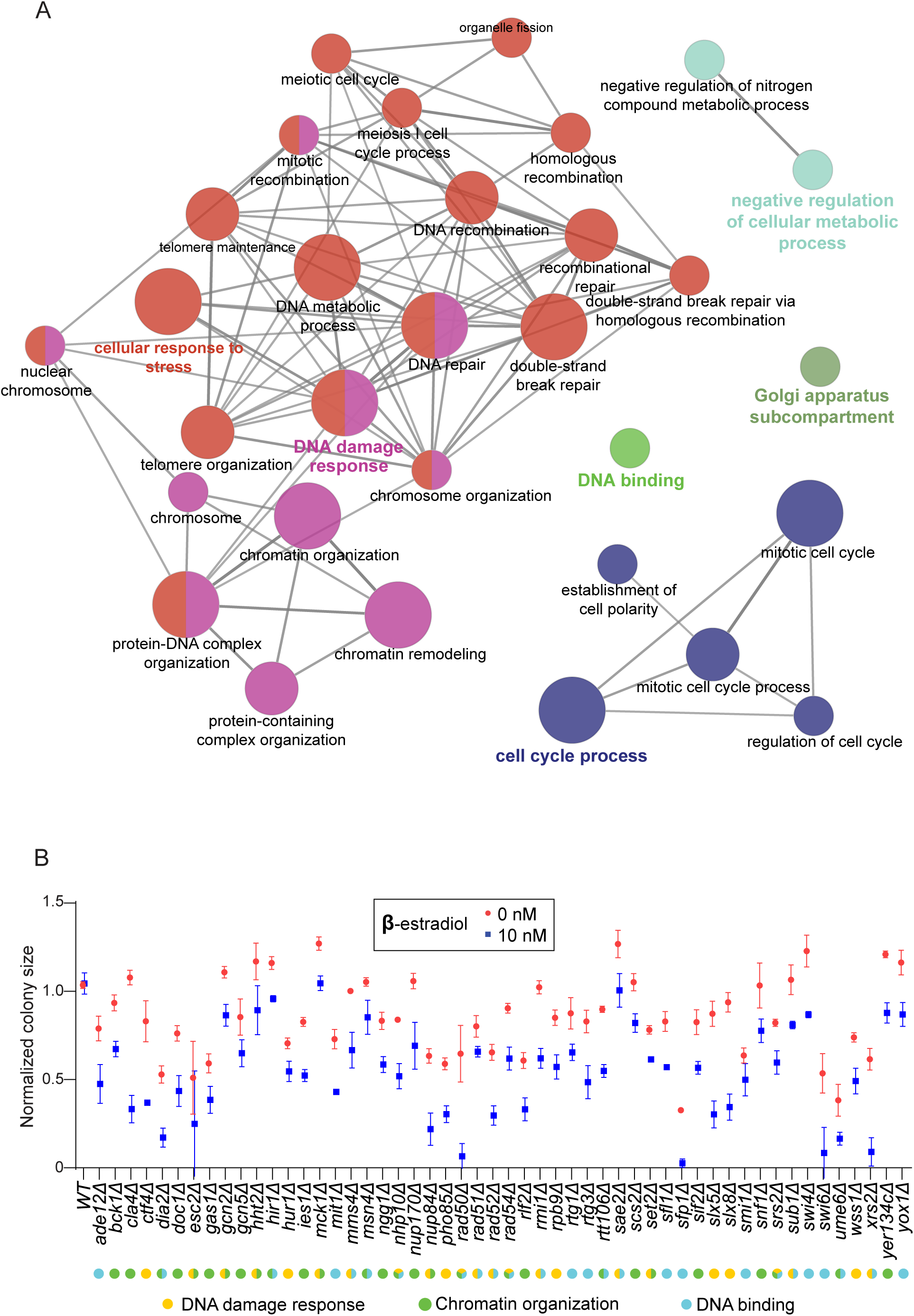
Results of SIM genetic interaction screen with MECP2-ref. (A) Functional enrichment for genetic interactions is shown where nodes indicate enriched GO terms and edges indicate statistically significant associations between terms. Node size and edge thickness indicate statistical significance (minimum p < 0.05, Bonferonni corrected). Highlighted labels indicate most significant terms within networks. Network generated using Cytoscape and the ClueGO plug-in. (B) Effect of MECP2-ref on yeast growth in mutants from the GO categories “DNA damage response”, “Chromatin organization” and “DNA binding” that were identified in the SIM screen. Normalized colony size indicates the size of the colonies on the array plates after accounting for the effect of MECP2-ref expression in WT yeast, in non-inducing (0 nM estradiol) and inducing (10 nM estradiol) conditions.

Looking more closely at these genes, we identified five out of the ten members of the *RAD52* epistasis group, *RAD50/51/52/54* and *XRS2* (*RAD55/57/59*, *RDH54* and *MRE11* were not identified) (Figure 3B). Mutants in this group are defective in the repair of DNA damage, maintenance of telomere length and meiotic and mitotic recombination (Symington 2002). We also identified three out of four of the subunits of the highly conserved MRX endo/exonuclease complex, which functions in repair of DNA double strand breaks, detection of damaged DNA, DNA damage checkpoint activation, telomerase recruitment and suppression of chromosomal defects (Syed and Tainer 2018). We identified Rad50, Xrs2 and Sae2, but not *MRE11* in our screen (Figure 3B). *RAD50* and *XRS2* were among the strongest interactions such that MECP2 expression reduced growth of the *rad50Δ* and *xrs2Δ* mutants to almost undetectable levels.

We also identified both subunits of the SBF transcription factor complex, Swi4 and Swi6. Swi6 is also a component of the MBF complex (MBF comprises Swi6 and Mbp1), which along with SBF functions to regulate transcription of the G1/S gene cluster as well as transcription of genes involved in DNA synthesis and repair (Bähler 2005). *swi6Δ* cells grow slowly, have a delayed G1/S transition and are sensitive to DNA damaging agents. Expression of MECP2 in the *swi6Δ* mutant severely reduced growth indicating a strong interaction. Additionally, we identified a strong interaction with the cyclin-dependent kinase *PHO85*, which regulates many cellular processes in response to stress and also plays an important role in G1/S transition via interaction with various G1 cyclins (Jiménez et al. 2013). *Δpho85* cells grow slowly, have increased mitotic chromosome loss and shortened telomeres. Taken together, these strong aggravating interactions with genes with central functions in DNA synthesis and repair, chromatin organization and G1/S cell cycle progression suggested MECP2 expression severely antagonized these processes which resulted in reduced fitness.

### Assessment of MECP2 variants in yeast

We decided to simply exploit the robust slow growth phenotype observed upon MECP2 expression in WT yeast to determine the effects of MECP2 variants, anticipating variants that interfered with MECP2 function would no longer reduce growth. We measured yeast growth in liquid culture for all 46 variants using the galactose-inducible 2-micron vector, because this vector produced the largest growth defect. We calculated exponential growth rates for WT yeast expressing MECP2 variants and using a linear mixed-effects model estimated the effects of the variants on MECP2 function (raw data is provided in Supplemental Data 4). Shown in Figure 4A is the model’s estimates of MECP2 function, arranged by variant amino acid position. Values were standardized to the range [0-1] where 0 represents the mean phenotype measured for an empty vector, and 1 represents the mean phenotype with the *MECP2-ref* construct. Hence, variants with values near to 1 functioned similarly to MECP2-ref, whereas values greater or less than 1 indicated increased or decreased function, respectively (error is plotted as 95% confidence intervals).

**Figure 4:**
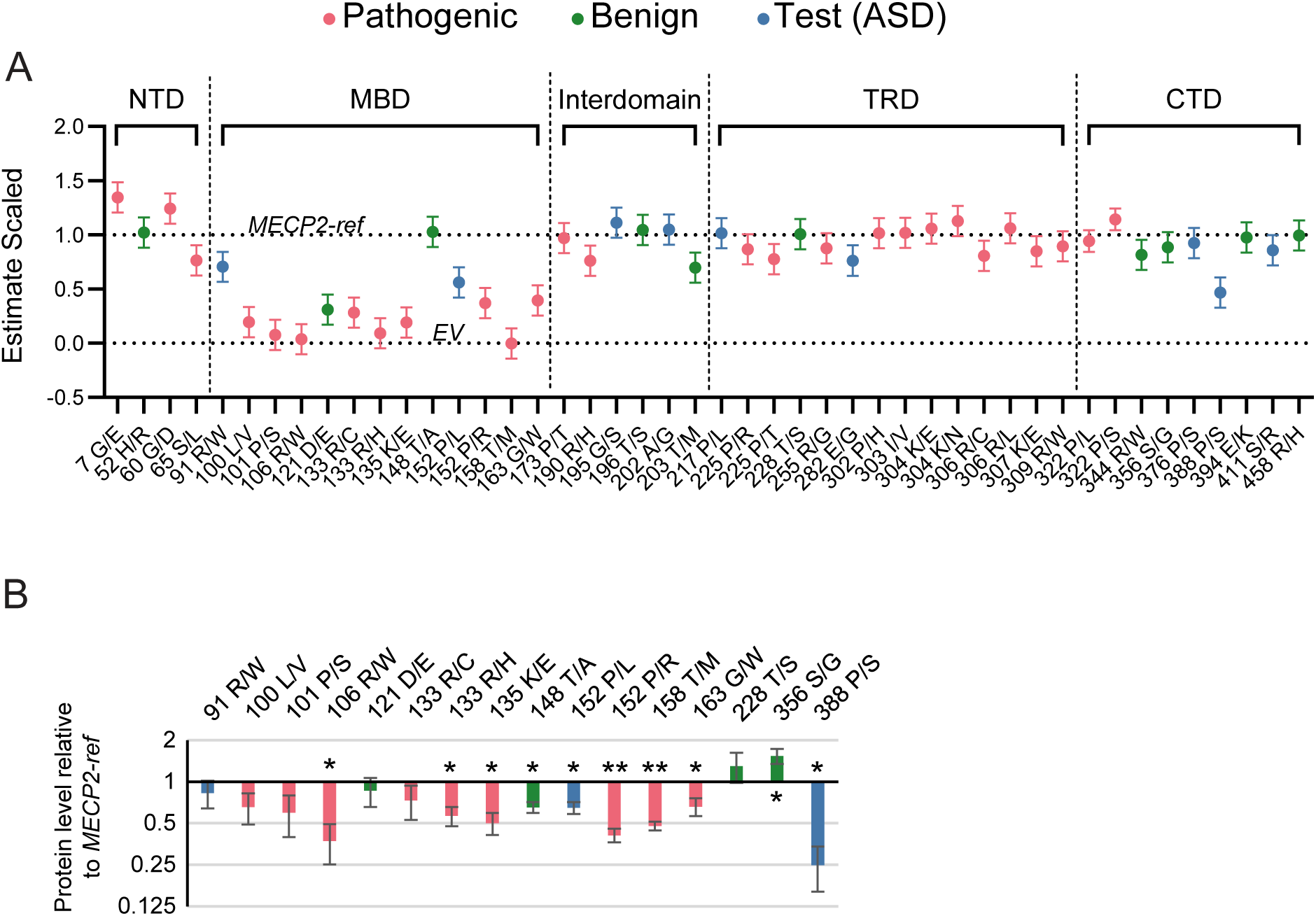
Analysis of 46 MECP2 variants using the yeast assay. (A) Estimated functional effects of MECP2 variants expressed in WT yeast. Data is normalized to MECP2-ref (=1) and empty vector (=0). Error bars indicate 95% confidence intervals. (B) Protein stability of MECP2 variants in WT yeast. Protein levels were normalized to MECP2-ref and plotted on a log scale. Error bars indicate S.D. (n = 3). * indicates p < 0.05, ** indicates p < 0.01. Colour coding same as in (A). All variants are located in MBD except T228S (TRD), S356G (CTD) and P388S (CTD).

Benign calibration variants had little effect on function as expected across all MECP2 domains. The one exception was the D121E variant which was nearly complete LoF, for which we have additional supporting evidence in fly leading to reclassification of this variant as pathogenic (see below). For the MBD domain, all seven pathogenic calibration variants were complete or near-complete LoF (we define a variant is complete LoF when its 95% confidence interval overlaps with 0). This supported that our assay was disease-relevant for variants in the MBD. L100V, R106W, R133C, and T158M have been previously reported to have reduced DNA binding *in vitro* and *in vivo* (Tillotson and Bird 2020), suggesting our assay reported on this function of the MBD. Residue R133 is part of the DNA interface and interacts directly with guanine of the mCpG island, and T158 is part of the Asx-ST motif that interacts with the DNA backbone (Tillotson and Bird 2020). The two test (ASD) variants in this domain, R91W and P152L, were found to have decreased function, although not as severe as the calibrating pathogenics. In contrast, calibrating pathogenics outside the MBD generally did not show decreased function. This indicated the yeast assay could not reliably detect altered function for variants outside the MBD.

### Protein level of variants in yeast

We measured protein levels for all variants in the MBD to determine if decreased stability/ increased turnover might be a factor in our assessment of function. We also included additional non-MBD calibrating benigns from the TRD (T228S) and the CTD (S356G) for comparison; and P388S because this variant showed the most reduced function of all non-MBD variants. MECP-2 protein levels from yeast extracts were measured on Western blots and normalized to MECP2-ref (Figure 4B and Supplemental Figure 1). For the MBD, 8 out of 13 MBD variants had reduced levels relative to MECP2-ref, while 5 were not significantly different from MECP2-ref. Variants with decreased protein levels ranged from ∼65% (T148A) to ∼37% (R106W) of MECP2-ref. The two calibrating benigns outside the MBD, T228S and S356G, did not show decreased protein levels, whereas the test (ASD) variant P388S had substantially reduced protein levels (∼ 25% of MECP2-ref).

Comparing protein levels to function of variants there appeared to be little correlation. Some variants had normal protein levels yet were non-functional (L100V, P101S, D121E, R133C). Other variants showed both decreased protein levels and decreased function (R106W, T158M, G163W, P388S). This suggested in some cases the variant affected MECP2 DNA binding directly leading to decreased function (e.g., R133C), whereas in other cases decreased function might also be a result of destabilization/ increased turnover (e.g., R106W). To investigate this in more detail we looked to variants outside the MBD, because these should not directly affect DNA binding. P388S was the only variant outside the MBD that showed moderate LoF (∼ 50%; Figure 3A) and we found it to have greatly reduced protein levels, whereas a variant in the same domain, S356G, was functional and had normal protein levels. The modest decrease in function of P388S relative to its low protein levels, suggested our functional assay was relatively insensitive to destabilization. The T148A variant, which had reduced protein levels (∼65% of MECP2-ref), but was completely functional in the assay, further supported this conclusion. Overall, these data suggested the yeast functional assay primarily reported on the intrinsic DNA binding ability of the MECP2 MBD.

### Using a fly wing vein assay for assessing MECP2 variant function

Expression of *MECP2* in *Drosophila* tissues has a wide range of phenotypes (Cukier et al. 2008; Vonhoff et al. 2012; Williams, White, et al. 2016; Williams, Mehler, et al. 2016). From these reports, we chose to assess MECP2 variants in developing wing tissues. This was because (1) the reported ectopic wing vein phenotype could be exploited here to provide a quantitative readout of MECP2 variant protein function, and (2) *MECP2* was shown to genetically interact with known MECP2 co-repressor complexes in the context of this specific phenotype (Cukier et al. 2008), suggesting the wing vein phenotype was relevant to MECP2 function in humans. These provided strong precedence for using this wing vein assay for assessing the function of ASD variants in MECP2.

We integrated *UAS-MECP2-ref* into the *attP40* locus on the *pGW.HA.attB* transgenic vector backbone to control for genome position effect and copy number (Bischof et al. 2007), to reduce variability in expression levels between integrants. In pilot experiments we tested a range of conditions including wing imaginal disc *GAL4* drivers, incubation temperatures and sex of progeny (as females have larger wings). We found that the ideal set of variables to produce a robust ectopic wing vein phenotype were the *A9-GAL4* driver in males (X-chromosome hyperactivation in male flies results in higher GAL4/UAS activity) raised at 29 °C (Figure 5, arrows). We also noted *MECP2-ref* expression resulted in smaller wings (Figure 5). Thus, we could proceed with assessing the effects of variants using this assay.

**Figure 5:**
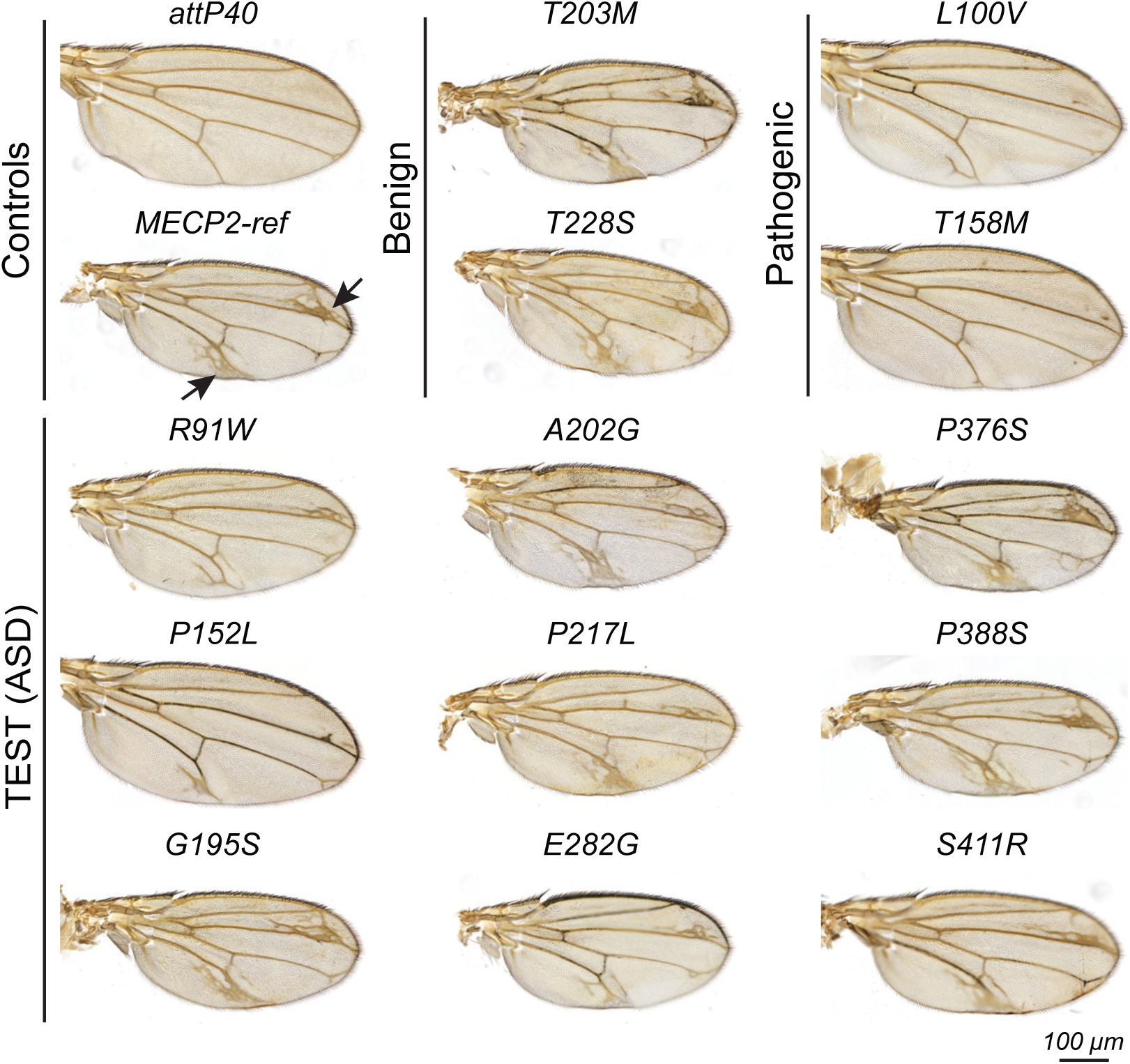
*Drosophila* wing phenotypes induced by MECP2 variants. Representative images of wings are shown expressing the indicated controls and variants. *attp40* – no MECP2, *MECP2-ref* – WT MECP. All images are the same scale. Arrows indicate ectopic vein tissue for *UAS-MECP2-ref* only.

### Assessment of 25 MECP2 variants using the wing vein assay

In total 25 transgenic lines were generated, including eight calibrating benign, eight calibrating pathogenic and nine test (ASD) variants. For all transgenic lines we passaged mating adults through three consecutive ‘assay’ vials. Adult progeny were then collected 3 days post-eclosion from each assay vial, and a single wing from a minimum of 10 adult male progeny per assay vial was removed and slide-mounted, for a total of 30 wings per variant. Mounted wings were scanned and total wing vein area / wing area was analyzed using a Labkit machine learning algorithm that semi-automated the quantitation of wing vein area (see Methods).

Next, we examined wing vein phenotypes induced by expression of all 25 MECP2 variants, with *attP40* and *MECP2-ref* controls in parallel. Representative images for controls, selected calibrating benign and pathogenic variants and all test (ASD) variants are shown in Figure 5 (also see Supplemental Figures 2 and 3 for representative images of all MECP2 variants and the masked area that was considered to contain vein tissue). Benign calibration variants all showed increased ectopic wing veins (T203M and T228S are shown) whereas calibrating pathogenics showed very little (L100V and T158M are shown), and test (ASD) variants had a range of outcomes (Figure 5). For example, P376S showed increased ectopic wing veins similar to *MECP2-ref* whereas P152L had few extra wing veins.

To quantify this phenotype, we trained two machine learning algorithms to quantify (1) the total area of the wing vein region and (2) the total area of the wing (see Supplemental Data 5 for details). Then using a linear mixed effects model these two values were used to calculate the estimated effect of variants relative to *MECP2-ref* and were normalized to *MECP2-ref* (at 1) and *attp40* (at 0) (Figure 6, error bars are 95 % confidence intervals). All benign calibration variants functioned similar to *MECP2-ref*, except for D121E. This variant was classified in ClinVar as benign by a single contributor, but we find both yeast and fly assays characterized it as a strong LoF, hence we predict D121E is likely pathogenic. All pathogenic calibration variants showed varying degrees of LoF, indicating the wing vein assay reported on disease relevant functions for MECP2. Most were strong LoF variants, except for P225R, which had reduced function but not a complete LoF. This successful calibration of the assay indicated clinically pathogenic and benign variants can be correctly classified in the fly model. We were particularly confident for variants in the MBD and TRD domains, because we tested multiple calibrating pathogenics in these domains, and both DNA and co-repressor interactions through these domains are validated in this assay. In the same experimental batch as the calibrating benign and pathogenic variants (controlling for food, temperature and humidity), we determined the wing vein phenotype for all nine test (ASD) variants. We found that both MBD variants R91W and P152L were LoF. The TRD variants P217L and E282G exhibited a partial LoF with data distributed similarly to the TRD pathogenic variant P225R. Finally, the Interdomain (G195S, A202G) and CTD variants (P376S, P388S and S411R) all exhibited normal function, similar to the calibrating benigns. Thus, we conclude that only test (ASD) variants in the MBD and TRD domains were LoF.

**Figure 6:**
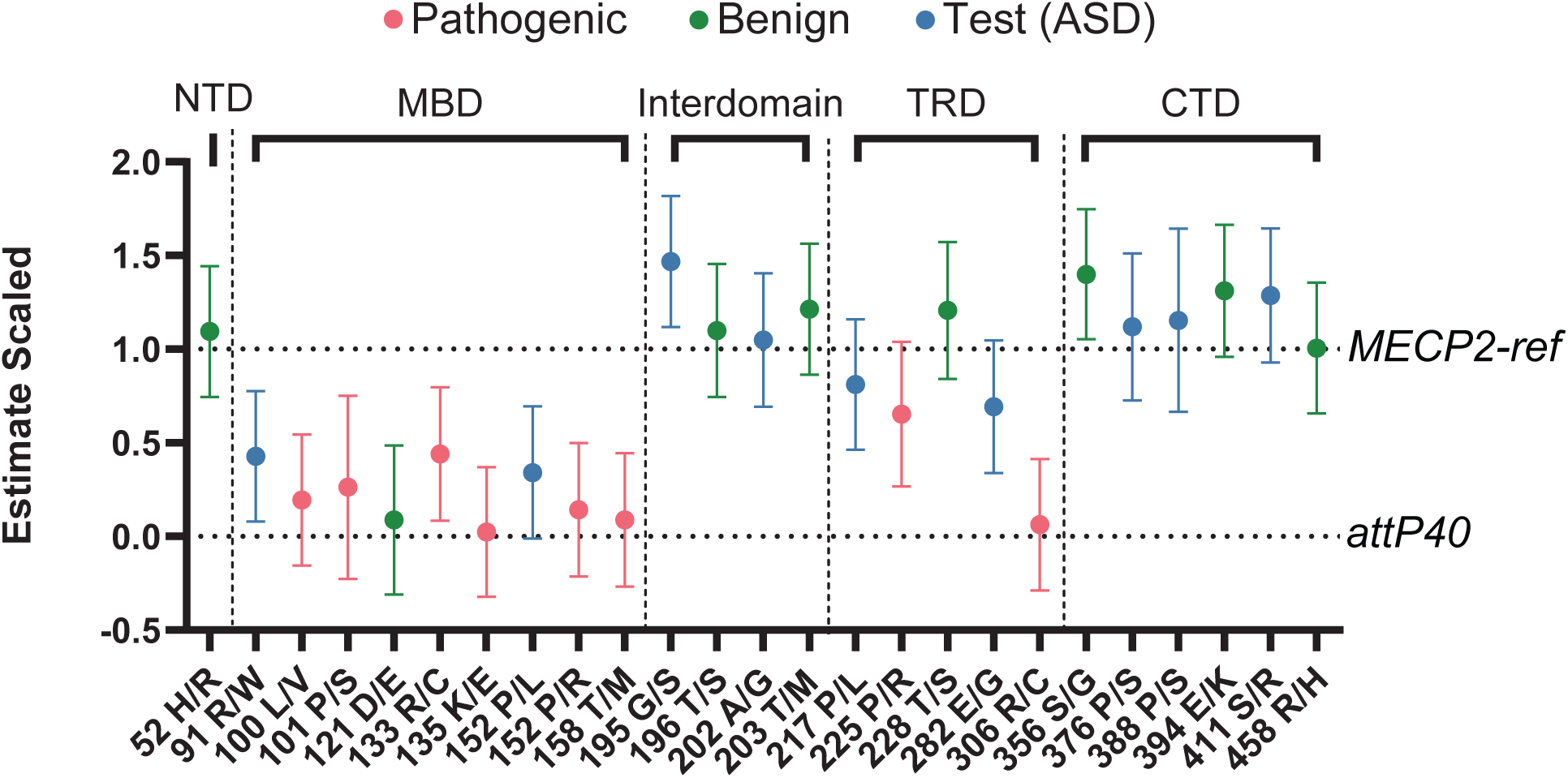
Analysis of MECP2 variant wing vein phenotypes. We show our quantitation of total wing vein per wing area for each genotype, normalized to *MECP2-ref* (WT MECP2) = 1 and *attp40* (no MECP2) = 0. Error bars indicate 95% confidence intervals.

Finally, we compared yeast and fly assay results for MBD variants only, because the yeast assay could only reliably measure MBD variants. We observed remarkable similarity in results between the two assays. Of particular interest was that both test (ASD) variants R91W and P152L appeared more functional in both yeast and fly assays than the pathogenic calibration variants. This suggested ASD variants may retain more DNA binding capacity than Rett variants, given these assays likely reported on the intrinsic ability of MECP2 to bind DNA.

## DISCUSSION

### Clinical classification of variants

Using our assays calibrated with variants classified in ClinVar as pathogenic and benign, we are able to provide functional assessments for nine test (ASD) variants, five of which have been found in cases of ASD without Rett (underlined): <u>R91W</u>, P152L, <u>G195S</u>, <u>A202G</u>, P217L, <u>E282G</u>, P376S, P388S, <u>S411R</u> (Figure 1). We found that R91W and P152L were LoF variants (yeast and fly) and we classify these as likely pathogenic. Our classification for P152L agrees with the updated pathogenic ClinVar classification and we re-classify R91W from a VUS in ClinVar. G195S had a modest gain of function (fly), hence we classify it as likely benign. Our likely pathogenic classification for G195S agrees with the updated benign ClinVar classification. A202G was found to be functional (fly). The ClinVar classification for this variant is conflicting (one benign and one VUS report) and we propose this variant be classified as likely benign.

P217L and E282G showed moderately reduced function (fly) consistent with their updated likely pathogenic ClinVar classifications. P376S, P388S and S411R were found to be functionally normal (fly). For P376S and P388S this is consistent with their updated benign classifications in ClinVar. However, since P388S appeared to be a destabilized protein in yeast we suggest P388S is worthy of further examination. We classify S411R as likely benign providing the first classification for this variant. Importantly, we provide functional evidence to classify two variants found in ASD but not Rett as likely pathogenic (R91W, E282G), suggesting certain mutations in MECP2 may cause ASD while others will cause Rett. Further, our data suggest variants found in ASD but not Rett may be less severe than Rett causing mutations, since the ASD-only variants we tested were more functional than Rett causing variants.

Finally, we provide evidence for reclassification of two variants, D121E and T148A, currently classified in ClinVar as benign and conflicting, respectively. D121E was LoF in both yeast and *Drosophila* assays, hence we propose D121E be reclassified as likely pathogenic. Interestingly, our data are consistent with *in vitro* studies showing a ∼7-fold reduction in the DNA binding capacity of the D121E variant (Free et al. 2001). This variant is neither in VariCarta nor RettBASE and it has one report in ClinVar, but this report lacks a description for how a benign classification was made. Interestingly, the computational predictors SIFT, PolyPhen and CADD all predict this variant is likely to be LoF (Supplemental Data 1), supporting our likely pathogenic classification. For T148A, the current ClinVar classification is conflicting. We find this variant to be functional (yeast), but to have modestly decreased protein levels (∼65% of MECP2-ref), suggesting it may have reduced stability. SIFT, PolyPhen and CADD all predict this variant to be damaging. We propose T148A be reclassified as likely benign, but suggest further characterization is warranted.

### Calibrating functional assays by protein domain

Often proteins contain multiple domains which contribute distinct functions to the protein. Our results indicate that when developing functional assays for human proteins in evolutionarily distant model organisms it is important to calibrate assays for each domain. This can be achieved through inclusion of calibration variants with known clinical/biochemical designations covering each domain. MEPC2 is a multi-domain protein and is largely unstructured outside the MBD domain (Ghosh et al. 2010). We initially focused on identifying calibrating variants in the MBD and TRD domains, as these domains harbour many of the clinically relevant variants.

The yeast assay only calibrated for the MBD and failed to calibrate for all other domains, indicating we could only assess functional effects of variants found within the MBD. A likely explanation is that the MECP2 MBD binds DNA in yeast without the need of accessory factors. Indeed, Brown *et al*., show this is likely to be the case (Brown et al. 2025 Mar 18). The TRD mediates interactions with co-repressor complexes, including Sin3A, N-Cor and REST, and Rett-causing mutations disrupt these interactions (Lyst et al. 2013). It is likely the assay failed to calibrate for TRD variants because these complexes are either not present in yeast (N-Cor and REST), or the TRD did not interact with the yeast ortholog (Sin3A/HDAC). The results of the MECP2 SIM screen also provided valuable insight into MECP2’s function in yeast. The strong aggravating genetic interactions we observed with genes involved in DNA damage response and DNA repair pathways (e.g., *RAD50*) suggest MECP2 expression either directly or indirectly results in DNA damage. These effects of MECP2 likely disrupted the cell cycle (supported by the interactions with cell cycle genes, e.g., *PHO85*) leading to overall decreased cell growth. Since variants in the MBD known to disrupt DNA binding failed to inhibit growth, this further suggested DNA binding by MECP2 contributed to defects in DNA and/or chromatin.

*Drosophila* was successful in stratifying pathogenic and benign calibration variants in both the MBD and TRD domains, indicating we can make high confidence predictions about test ASD variants located in these domains. For variants in the MBD, it is likely these disrupt DNA binding, although DNA binding by MECP2 has yet to be demonstrated in *Drosophila*. For the TRD, our data suggest MECP2 is able to interact with *Drosophila* orthologs of co-repressor complexes. First, R306C prevents interaction with co-repressors and was complete LoF in our assay (Lyst et al. 2013). Second, co-repressor complex mutants suppress phenotypes resulting from MECP2 over-expression in the *Drosophila* wing (Cukier et al. 2008). For example, mutation of the Polycomb group co-repressor ortholog of Asxl1, *Asx*, prevents ectopic wing vein growth resulting from MECP2 expression (Cukier et al. 2008). For the Interdomain and CTD domain, testing additional pathogenic variants in these domains would increase confidence in our predictions for test (ASD) variants within these domains. Thus, when developing functional assays in surrogate model systems with considerable evolutionary distance from humans, it is important to consider calibrating assays for each domain before making conclusions about the effects of test variants.

### MECP2 protein stability

The stability of a handful of MECP2 variants has been studied both using *in vitro* assays and *in vivo* mouse models (Kucukkal et al. 2015; Yang et al. 2016; Sperlazza et al. 2017; Tillotson and Bird 2020). We were able to compare seven variants located in the MBD between our yeast results and these studies (Supplemental Table 1). R106W, P152R and T158M are all destabilized in mouse models and *in vitro* systems, and based on protein levels in yeast were predicted to be also destabilized in yeast also. R133C is found to be stable in *in vitro* systems and is predicted to be stable in yeast, but is destabilized in mouse. L100V and P101S are destabilized *in vitro*, but are predicted to be stable in yeast. R133H is stable *in vitro*, but is predicted to be destabilized in yeast. Hence, the discrepancies between these data emphasize the value of comparing variant function across multiple species and assay systems. MECP2 turnover in cells is regulated by ubiquitylation by the E3 ligase RNF4 (Wang 2014), indicating it may be an *in vivo*-specific modifier of variant stability. Interestingly, we identified the yeast orthologs of RNF4, *SLX5* and *SLX8*, as aggravating genetic interactions in our SIM screen (Figure 3B), suggesting MECP2 levels may also be regulated by this E3 ligase in yeast. Thus, regulated turnover *in vivo* is likely an important factor contributing to MECP2 variant function and may vary across assay systems. Interaction of MECP2 with DNA and co-repressors also influences MECP2 intrinsic stability (Ghosh et al. 2010), hence differences in these interactions could be another factor contributing to differences in MECP2 variant function/stability between assay systems.

### MECP2 and DNA methylation

It is noteworthy that all pathogenic calibration variants in the MBD were LoF in our functional assays despite there being no, or extremely low levels of DNA cytosine methylation in yeast and *Drosophila*, respectively. Our data correlate well with Brown *et al*. (Brown et al. 2025 Mar 18), showing that MECP2 binds the unmethylated yeast genome and mediates widespread transcriptional suppression. This raises the possibility that ASD and Rett MECP2 variants may not be specifically deficient in their ability to bind methylated DNA, but may be generally deficient in binding to unmethylated DNA and/or chromatin. It is challenging to specify differential impacts of specific Rett variants on methyl versus non-methyl DNA binding in models where DNA methylation is present. We propose this may have obscured the impact that pathogenic variants have on non-methylated DNA binding and its importance in healthy and disease states, and we propose that the yeast and fly models offer valuable platforms to further examine these mechanisms.

## DATA AVAILABILITY STATEMENT

Strains and plasmids are available upon request. The authors affirm that all data necessary for confirming the conclusions of the article are present within the article, figures, tables and supplemental data files.

## ACKNOWLEDGEMENTS

We thank Dr. Janel Kopp, UBC for access to the Panoramic Midi II digital slide scanner. Susan Lindquist for the generous gift of the *pAG426GAL-ccdB* vector. Stocks obtained from the Bloomington *Drosophila* Stock Center (NIH P40OD018537) were used in this study. We thank *Drosophila* Genomics Resource Center, supported by NIH grant 2P40OD010949 for the *esc* cDNA. We used FlyBase (releases 2020 through 2024) for general information on genes, phenotypes and stocks. We would like to thank Gino Laberge at Genome ProLab for the kind gift of balancer stocks and also their transgenic services (Genome ProLab Inc, Quebec, Canada). We thank Libby Natola and Guillaume Poirier-Morency (Pavlidis Lab) for contributions to the variant annotation pipeline. This work was funded by an operating grant from The Simons Foundation Autism Research Initiative (2021 Genomics of ASD: Pathway to Genetic Therapies: AWD-021951. SIMOFOUN 2021).

## AUTHOR CONTRIBUTIONS

DWA, CJRL, BPY, SMG, JTC, MD, SR, MD, PP contributed to conception of the work.

DWA, CJRL, GM, EC, BPY, SI, TL, JL, JBL, SMG, BB, MD, JS, SR, PP contributed to design of the work.

DWA, CJRL, GM, EC, BPY, JBL, LM, JS, JBL, SR, PP contributed to acquisition, analysis, or interpretation of data.

DWA, SR, PP, GM, EC, JBL, MD, KKC, BB, BW contributed to creation of new software used.

DWA, CJRL, JS, BPY, JTC, JBL, GM, EC, LM, JL, JBL, JTC, SR, PP contributed to drafting the paper or substantively revised it.

## SUPPLEMENTAL FIGURE CAPTIONS

**Supplemental Figure 1:**
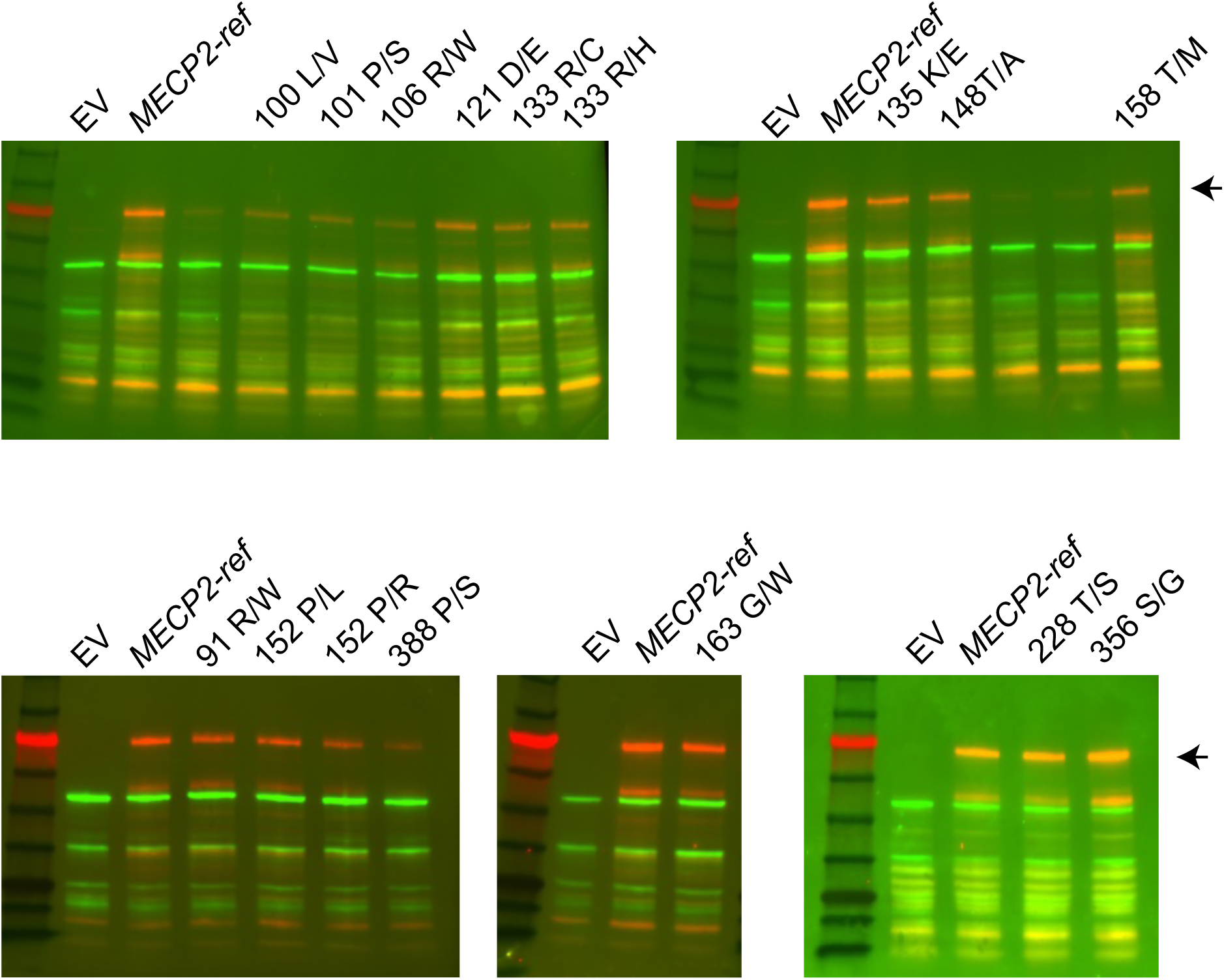
Representative MECP2 Western blots. Arrow indicates the full-length MECP2 protein. Antibody labelling is as follows: red - anti-MECP2, green - anti-PGK1.

**Supplemental Figure 2:**
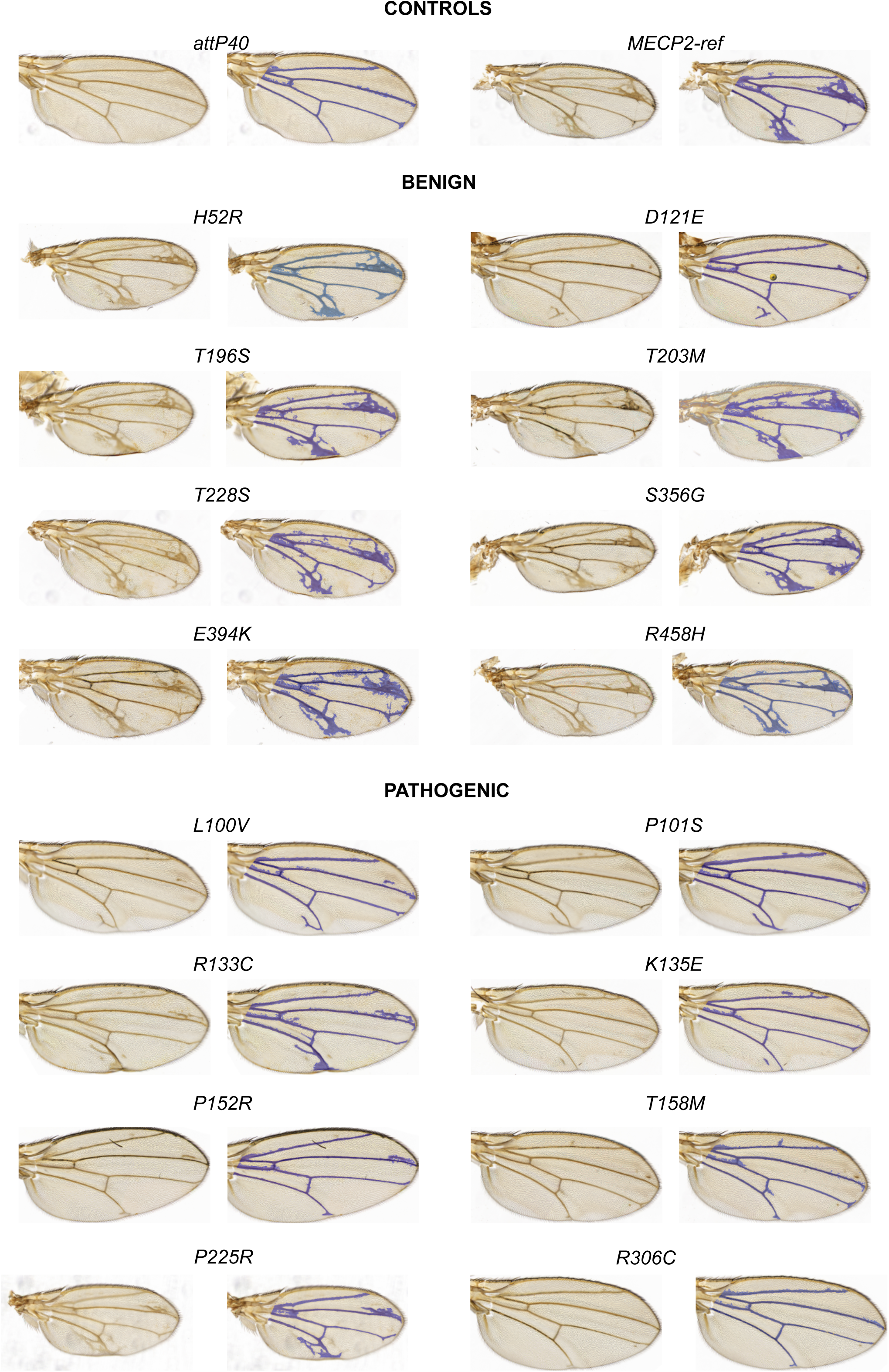
Representative wing phenotypes for control, benign and pathogenic MECP2 calibration variants tested in *Drosophila*. For each variant, raw images are on the left and the masked image used to quantitate total wing vein per wing is on the right.

**Supplemental Figure 3:**
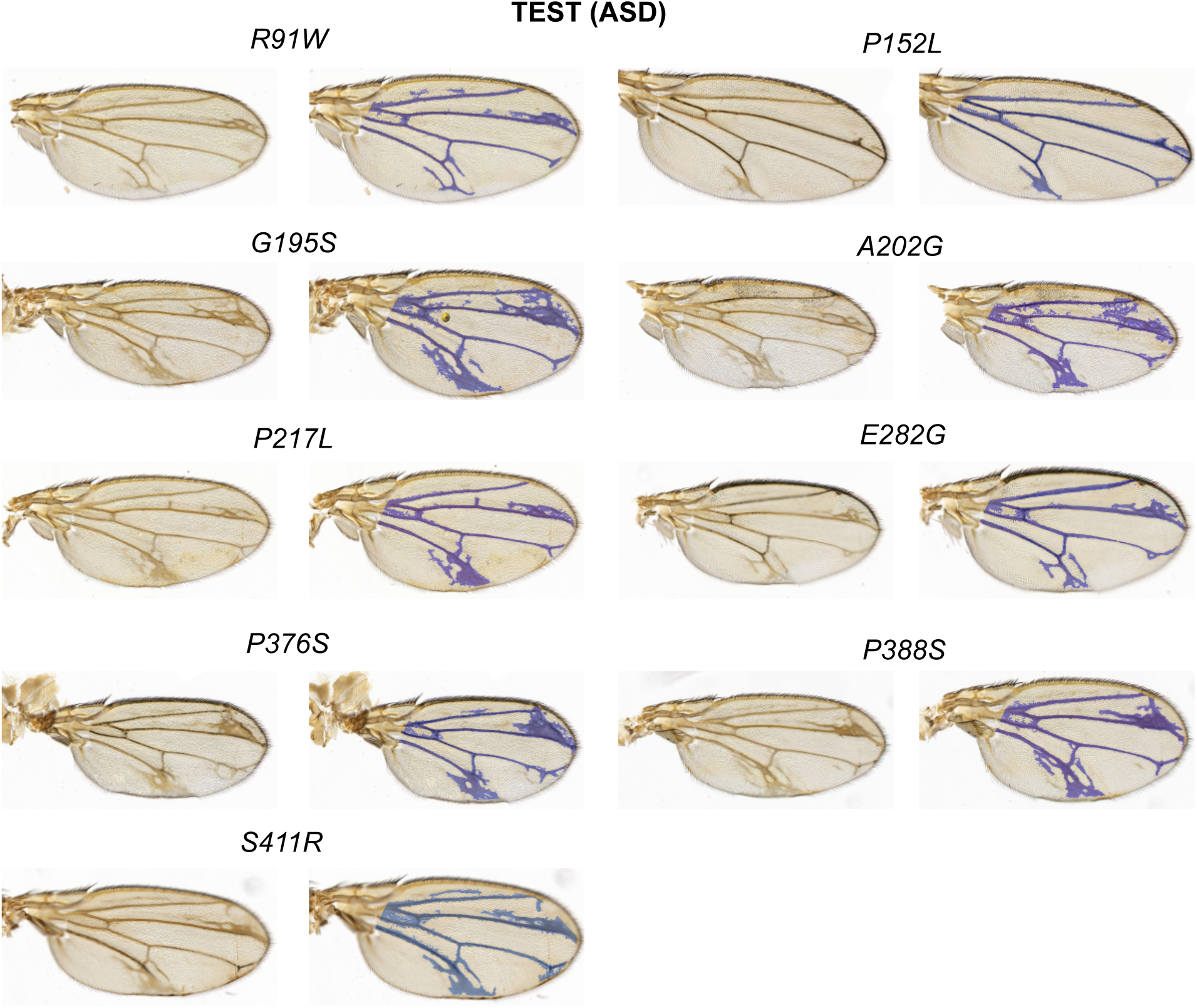
Representative wing phenotypes for test (ASD) MECP2 variants tested in *Drosophila*. For each variant, raw images are on the left and the masked image used to quantitate total wing vein per wing is on the right.

## SUPPLEMENTAL METHODS CAPTIONS

**Supplemental Methods 1:** Detailed outline of wing vein quantitation methods.

**Supplemental Methods 2:** Yeast statistical model R script.

**Supplemental Methods 3:** Fly statistical model R script.

## SUPPLEMENTAL TABLE CAPTIONS

**Supplemental Table 1:** MECP2 protein stability summary table.

| Variant | Yeast*<br>(this study) | Yang <sup>1</sup><br>(in vitro, WT = 1) | Kucukkal <sup>2</sup><br>(in vitro) ($\Delta\Delta G$) | Sperlazza <sup>3</sup><br>(NMR)<br>(+ slight, ++ more destabilized) | Tillotson <sup>4</sup><br>(mouse models, WT = 1) |
| --- | --- | --- | --- | --- | --- |
| R91W | 1 |  |  |  |  |
| L100V | 1 | 0.5 | +1.25 |  |  |
| P101S | 1 |  |  | + |  |
| R106W | 0.37 | 1 | +0.1 | ++ | 0.1 (TAVI) |
| D121E | 1 |  |  |  |  |
| R133C | 1 | 1 | +0.4 |  | 0.6 (eGFP) |
| R133H | 0.56 | 1 | +0.25 |  |  |
| K135E | 0.50 |  |  |  |  |
| T148A | 0.65 |  |  |  |  |
| P152L | 0.65 |  |  |  |  |
| P152R | 0.41 | 0.25 | +1.5 |  |  |
| T158M | 0.48 | 1 | +0.25 | + | 0.3 (eGFP), 0.25 (TAVI) |
| G163W | 0.66 |  |  |  |  |
| P225R |  |  |  |  | 0.22 |
| T228S | 1 |  |  |  |  |
| R306C |  |  |  |  | 1.0 |
| P322L |  |  |  |  | 0.03 |
| S356G | 1.54 |  |  |  |  |
| P388S | 0.25 |  |  |  |  |

\* Values not significantly different than MECP2-Ref are set to 1, all others are significant (p<0.05)

## SUPPLEMENTAL DATA CAPTIONS

**Supplemental Data 1:** Detailed information for each variant, primers used in cloning, the molecular genetic reagents made for each organism, and the *Drosophila* stocks generated.

**Supplemental Data 2:** Yeast SIM screen raw data.

**Supplemental Data 3:** GO analysis results for yeast SIM screen listing genes in each GO term.

**Supplemental Data 4:** Raw data for yeast variant assays used in statistical model.

**Supplemental Data 5:** Raw data for *Drosophila* variant assays used in statistical model.

